# Spatiotemporal Dynamics of Stemphylium Leaf Blight and Potential Inoculum Sources in New York Onion Fields

**DOI:** 10.1101/2021.07.30.454532

**Authors:** Frank Hay, Daniel W. Heck, Audrey Klein, Sandeep Sharma, Christy Hoepting, Sarah J. Pethybridge

**Affiliations:** Plant Pathology & Plant-Microbe Biology Section, School of Integrative Plant Science, Cornell AgriTech, Cornell University, Geneva, NY 14456; Cornell Vegetable Program, Cornell Cooperative Extension, Albion, NY 14424

## Abstract

Stemphylium leaf blight (SLB) caused by *Stemphylium vesicarium* is the dominant foliar disease affecting large-scale onion production in New York. The disease is managed by fungicides, but control failures are prevalent and recently attributed to fungicide resistance. Little is known of the relative role of inoculum sources in initiation and spread of SLB epidemics. The spatial and spatiotemporal dynamics of SLB epidemics in six onion fields were evaluated along linear transects in 2017 and 2018. Average SLB incidence increased from 0 to 100% throughout the cropping seasons with an average final lesion length of 28.3 cm. Disease progress was typical of a polycyclic epidemic and the logistic model provided the best fit to 83.3% of the datasets. Spatial patterns were better described by the beta-binomial than binomial distribution in half of the datasets and random patterns were more frequently observed by the index of dispersion. Geostatistical analyses of spatial pattern also found a low frequency of datasets with aggregation. Spatiotemporal analysis of epidemics detected that the aggregation was influenced by disease incidence. However, diseased units were not associated with the previous time period according to the spatiotemporal association function of *SADIE*. Variable spatial patterns suggested mixed inoculum sources dependent upon location, and likely an external inoculum source at the sampling scale used in this study. Plate testing of 28 commercially available organic onion seedlots from 2017 and 2018 did not detect *S. vesicarium*. This finding suggests that although *S. vesicarium* has been reported as seed transmitted, this is unlikely to be a significant inoculum source in commercially available organic seed lots and even less so in fungicide-treated seed used to establish conventional fields. A small-plot replicated trial was also conducted in each of two years to quantify the effect of *S. vesicarium*-infested onion residue on SLB epidemics in a field isolated from other onion fields. SLB incidence was significantly reduced in plots without residue compared to those in which residue remained on the soil surface. Burial of infested residue also significantly reduced epidemic progress in one year. The effect of infested onion residue on SLB epidemics in the subsequent onion crop suggests rotation or residue management may have a substantial effect on epidemics. However, the presence of an inoculum source external to fields in onion production regions as indicated by a lack of spatial aggregation may reduce the efficacy of in-field management techniques.

---

New York (NY) is the fifth largest producer of onions in the USA, producing 142,246 t from 2,832 ha (USDA NASS 2017). Large-scale onion production occurs in localized regions of mostly muck soil, a type of histosol high in decomposed organic matter. In these regions, onion is the predominant crop and rotation occurs only infrequently to other root vegetables such as potato, table beet, and carrot. Crops are established by transplants or seed from late March to early June, and harvests extends from mid-July to late October (Leach et al. 2018). Onion production is affected by annual epidemics of foliar fungal diseases including Botrytis leaf blight (*Botrytis squamosa*), purple blotch (*Alternaria porri*), downy mildew (*Peronospora destructor*), and Stemphylium leaf blight (SLB; *Stemphylium vesicarium*) (Brewster 2008). SLB was first reported on onion in the United States in Texas (Miller et al. 1978) and was subsequently reported in NY in 1985 (Shishkoff and Lorbeer 1987, 1989). A major epidemic of SLB occurred in NY in 1990, with SLB present in almost all onion fields surveyed across NY, causing severe foliar dieback in some fields (Lorbeer 1993). However, SLB occurred only sporadically in onion in 1991 and 1992, and was rare in 1993 (Lorbeer 1993). Subsequently SLB was considered a minor component of the foliar disease complex affecting onion in NY but has become predominant since 2010 (Hay et al. 2019). Crop loss from SLB manifests as small bulbs of poor quality associated with premature loss of foliage and the inability to properly lodge (Brewster 2008). For example, when the disease is severe, SLB has been associated with up to 74% failure to lodge in onion plants (Hoepting 2017). Foliar disease management aims to ensure photosynthetic area is maximized throughout the growing season to achieve optimal bulb size and quality. In a NY field trial, selected fungicides resulted in a 28.7% increase in the number of jumbo grade bulbs, attributed to enhanced foliar health from improved SLB control (Hoepting 2018). The premature leaf senescence associated with SLB is also associated with an increased incidence of bacterial bulb rot and deleteriously impacts storage quality (Paibomesai et al. 2012). In many crops in NY maleic hydrazide is applied near harvest to reduce the sprouting of bulbs in storage. However, optimal uptake of the sprout inhibitor requires five to seven green leaves per plant. If insufficient maleic hydrazide is taken up due to poor foliar health from SLB and other diseases, bulbs will sprout in storage and cannot be sold (Brewster 2008; Isenberg et al. 1974).

SLB lesions are initially water-soaked and become pale brown to tan in color. They usually appear first on the older leaves and are typically oval shaped but may become spindle to ovate-elongate with age. Profuse sporulation of *S. vesicarium* occurs on the lesions and other necrotic tissue leading to a gray to dark olive brown appearance. Colonization of leaves by *S. vesicarium* early in the season often begins in leaf tips that have become necrotic due to other factors, e.g., herbicide damage. Leaf tips often become colonized by a range of fungi including *Alternaria* spp. and *S. vesicarium* and this symptom is commonly referred to as ‘dirty tips’. Conidia of *S. vesicarium* formed on leaf tips may provide secondary inoculum, or the necrosis may continue to progress down the leaf, often asymmetrically (Miller and Schwartz 2008; Hay et al. 2019).

Relatively little is known of how *S. vesicarium* is introduced and the relative contribution of potential overwintering inoculum sources for SLB epidemics in onion fields. *S. vesicarium* is known to be seed transmissible in onion (Aveling 1993; Aveling et al. 1993) and infested seed was hypothesized as a primary inoculum source in SLB epidemics in NY onion fields in 1985 and 1990 (Lorbeer 1993), but not conclusively demonstrated. Transplants may also have potential to act as a source of SLB. This may occur either by introducing diseased transplants from outside NY, or by providing an early crop upon which SLB can establish, build up, and subsequently provide inoculum for direct seeded crops nearby. Volunteer onions from previous seasons may also present a source of inoculum, especially in NY muck regions where onion fields are in close proximity and there is often an absence of crop rotation. *S. vesicarium* has a broad host range encompassing common annual and perennial crop and weed species from several families (Farr and Rossman 2020), which may contribute as alternative hosts and green bridges to initiate SLB epidemics in onion fields. For example, *S. vesicarium* also causes economically important diseases in other Alliums, such as leek and garlic (Miller and Schwartz 2008), and in soybean (Darrag et al. 1982; Lamprecht et al. 1984), alfalfa (Lowe et al. 1987), pear (Kӧhl et al. 2009; Llorente and Montesinos 2006; Rossi et al. 2005a, 2005b, 2008, 2009), parsley (Koike et al. 2013), tomato (Trinetta et al. 2013), radish (Belisario et al. 2018), and asparagus (Bohlen-Janssen et al. 2018; Foster et al. 2019). Host-strain specificity within *S. vesicarium* populations has also been reported (Kӧhl et al. 2009). Singh et al. (1999) reported that *S. vesicarium* isolates from asparagus were pathogenic to Japanese pear but not European pear. Moreover, *S. vesicarium* isolates from onion and asparagus were not pathogenic to pear (Köhl et al. 2009). However, in other studies, *S. vesicarium* isolates from asparagus, onion, and garlic were pathogenic to all three crops (Basallote-Ureba et al. 1999) and isolates from onion were pathogenic to asparagus (Foster et al. 2019). Crops, weeds, and potentially inter-crop species (e.g., barley) may also serve as alternative pathogen hosts. *S. vesicarium* can act as a saprophyte and colonize and sporulate on dead grass species (Lamprecht et al. 1984; Kohl et al. 2009; Rossi et al. 2005b).

*S. vesicarium* overwinters on infested residue as mycelia or the teleomorph, *Pleospora allii* (Simmons 1969; Inderbitzin et al. 2005). Given that onion production on muck soil is largely devoid of crop rotation, infested onion residue in soil is likely to be a dominant inoculum source between seasons. *S. vesicarium* is homothallic and both mating types have been identified in the genomes of NY isolates (Inderbitzin et al. 2005; Sharma et al. 2020). In NY, pseudothecia occur commonly in diseased onion leaves at harvest, and while there have been no studies of ascospore release in the field in NY, ascospores have been observed to form within the pseudothecia upon cool storage of infested onion leaves for several weeks (F. S. Hay, *unpublished data*). In Ontario Canada, *P. allii* ascospores have been found in spore traps in early spring before the onion crop is planted (Gossen et al. 2021). This suggests that ascospores may initially colonize alternative hosts or volunteer onions before conidial spread through multiple infection cycles within the onion growing season. The role of the teleomorph and other inoculum sources in the epidemiology of SLB in NY onion fields is a critical knowledge gap.

Quantifying the spatiotemporal characteristics of epidemics provides valuable information upon which to develop hypotheses surrounding inoculum sources, pathogen dissemination and epidemic progress (e.g., Madden 1980; Pethybridge et al. 2005; Cieniewicz et al. 2018; Gigot et al. 2017; Heck et al. 2021), which is lacking for SLB of onion. Spatiotemporal characteristics of SLB epidemics are also vital for evaluating crop loss (Madden and Hughes 1995; Madden et al. 2018), experimental design (Madden and Hughes 1995; Madden et al. 2018), the design of sampling methods (Turechek and Madden 1999, 2004; Turechek et al. 2011; Jones et al. 2011; Heck et al. 2021; Madden et al. 2018) and management strategies (Ristaino and Gumpertz 2000; Madden et al. 2007).

The objectives of this study were to: (i) to characterize and quantify the spatial and spatiotemporal dynamic of SLB epidemics in NY onion fields; (ii) assess the potential for seed to be infested with *S. vesicarium*; and (iii) to examine the role of infested onion residue on SLB epidemics in the subsequent onion crop. Filling in the epidemiological gaps surrounding the spatial distribution, dispersal and survival of the pathogen can help to develop hypotheses to evaluate efficient management strategies.

## Materials and Methods

### Spatial and temporal analyses of SLB epidemics

#### Data collection

The incidence and severity of SLB was assessed in each of three commercial, conventional large-scale onion fields in Elba, NY (43°08’ N; 78°07’ W) during the 2017 and 2018 cropping seasons. This region is a centralized area of onion production on muck soil. Each field varied between ∼ 5 and 15 ha in size. Sampling was conducted in a cluster sampling approach (Hughes et al. 1996). Three onion rows were arbitrarily selected within each field and operationally defined as linear transects. One of the transects was located at the field edge and the other two initiated at 25 and 50 m from the field edge. In 2017, samples were collected on 26 June, 7 and 27 July, and 7 and 22 August. In 2018, samples were collected on 20 June, 3 and 18 July, and 23 August. At each time, one fully expanded outer onion leaf was randomly collected from each of five plants at 3.05-m intervals at each of 31 locations per transect. The three transects within fields at each evaluation time were considered a dataset, the sampling location within transects as a sampling unit, and leaves as individuals. Overall, three fields were assessed each year, which corresponded to six datasets including each of 93 sampling units (*N* = 93) and 5 individual leaves (*n* = 5) per sampling unit, evaluated at each time. Leaves were returned to the laboratory and stored at 4°C for up to 5 days before disease intensity evaluation.

#### Disease intensity

Leaves were assessed for disease incidence (presence/absence) and then further incubated individually at 20°C in plastic bags with moist towel to encourage fungal sporulation. After 7 days, leaves were assessed for the presence of *S. vesicarium* conidia and/or *P. allii* pseudothecia using a stereo microscope (×60 magnification). Areas of the onion leaf with *S. vesicarium* sporulation were marked on the plastic bag and expressed as lesion size (longitudinal length of the diseased leaf portions accompanied by *S. vesicarium* sporulation). Lesion size was therefore used to represent disease severity. Ten isolations were conducted from putative SLB lesions on onion leaves per field at each sampling time to confirm the association with *S. vesicarium* as part of a previously reported study (Hay et al. 2019).

SLB incidence (*y*) in proportion was expressed for each sampling unit as *y_i_* = *x_i_*/*n*, where *x* was the number of diseased leaves in the *i*^th^ sampling unit (*i* = 1, 2, 3, …, *N*) containing *n* (= 5) leaves. Thus, the overall incidence for each dataset was calculated as *y* =∑*x*/(*N*×*n*), where *N* (= 93) is the total number of sampling units in each dataset. The same formula was used to summarize lesion size, with a modification replacing the number of diseased individuals with the total lesion length on each leaf.

#### Disease progress

The monomolecular, logistic and Gompertz models were fitted to SLB incidence over time by nonlinear regression using the nlsLM function from MINPACK.LM package (Elzhov et al. 2016). The model of best fit was selected based on the lower root mean square error (RMSE), independence and homogeneity of variances, and larger coefficient of determination (*R^2^*). The model parameters: progress rate (*r*); disease incidence at epidemic initiation (*y_0_*); time (days) to when disease incidence reached 50% (*y_50_*); incidence at the last assessment (*y_max_*); and the relative area under disease progress stairs (rAUDPS; Simko and Piepho 2012) were used for comparisons between cropping seasons. The rAUDPS was also calculated for lesion length. Comparisons between the cropping seasons were performed by a *t*-test with a Welch’s correction assuming unequal variances and sample sizes (Welch 1947). Analyses were performed in the R software v. 3.6 (R Core Team 2020).

#### Weather data

Daily recordings of temperature, relative humidity, dew point, and precipitation predicted for the centroid of the muck region on Elba, NY were obtained by the NASAPOWER package in R (Sparks 2018).

#### Spatial analyses

Two approaches, point-pattern and geostatistical-based, were used to characterize the heterogeneity of diseased individuals in the datasets. Point-pattern approaches did not account for the spatial location of diseased units and were used to characterize the spatial heterogeneity at the sampling unit and below. The analyses were performed by fitting the data to distributions and calculating the index of dispersion.

First, the binomial and beta-binomial distributions were fitted and compared to characterize the spatial pattern of SLB incidence (Madden and Hughes 1995; Turechek et al. 2011). A best fit to the binomial distribution is characteristic of a random pattern, while the beta-binomial provides a best fit to aggregated patterns (Madden et al. 2007). The parameters of the binomial, π, and beta-binomial distributions, *p* and *θ*, were estimated by maximum likelihood (Smith 1983; Sparks et al. 2008) and the fit to distributions determined by the χ^2^ goodness-of-fit test. A log-likelihood ratio test (LRS) was performed to understand when the beta-binomial provided a better fit to the observed data than the binomial distribution (Madden and Hughes 1994, 1995; Madden et al. 2018). The analyses were performed using the FIT_TWO_DISTR function from the EPIPHY package in R (Gigot 2018).

The index of dispersion (*D*) was calculated as a ratio between the observed and expected variances on the assumption of a binomial distribution (Madden et al. 2007). When *D* = 1, the data is likely to have a random pattern. When *D* < 1, the spatial pattern is likely to be regular, and if *D* > 1, the spatial pattern is aggregated. *D* has a χ^2^ distribution under the null hypothesis of randomness (Madden and Hughes 1995). Analyses were performed using the AGG_INDEX function within the EPIPHY package in R.

The geostatistical-based approach considers the spatial location of sampling units in the analyses. Three methods: runs analyses, spatial autocorrelation, and Spatial Analyses by Distance IndicEs (*SADIE*), were used to evaluate the spatial distribution of SLB incidence at the level of sampling unit and above.

Ordinary and median runs analyses were used to characterize patterns among sampling units in rows (Madden et al. 1982). For ordinary runs, the sampling units were classified as 1 or 0, if at least one diseased leaf or only asymptomatic leaves were present in the sampling units, respectively. For median runs, the sampling units were classified as 1 or 0 if incidence was above or below the median incidence of the dataset, respectively. The observed number of runs was compared with the expected number of runs by a Z-statistic assuming a null hypothesis of randomness (Gibbons 1976; Madden et al. 1982). An appropriate function was developed in R to perform the ordinary and median runs analyses based on formulas available in literature (Gibbons 1976; Madden et al. 1982).

Spatial autocorrelation statistics were used to quantify the degree of spatial dependence between regionalized disease incidence (Chilès and Delfiner 2012; Cressie 1989; Reynolds and Madden 1988). The proportion of diseased leaves in each sampling unit was transformed using the logit transformation *ln*(*y*/(1 – *y)*), where *y* was primarily corrected by the Haldane factor, *y* = (*x* + 0.5)/(*n* +1), for which *x* is the number of diseased leaves and *n* is the total number of leaves in a sampling unit (Reynolds and Madden 1988). Spatial autocorrelation coefficient functions (ACFs) were calculated up to lag 15 with a confidence interval of 95% for each spatial lag (Turechek and Madden 1999; Turechek and Mahaffee 2004). Analysis was performed using the ACF function within the STATS package in R.

*SADIE* uses the number of diseased individuals and the location of each sampling unit to quantify the distance to regularity, *D_r_*, defined as the minimum distance of shifts until all sampling units contain the same number of diseased individuals (Perry 1995, 1998; Perry et al. 1999). The index of aggregation (*I_a_*) is the ratio between the observed distance and the expected average distance to regularity based on randomizations, *I_a_* = *D_r_*/*E_r_* (Perry et al. 1999). If *I_a_* = 1, the random pattern is assumed, while *I_a_* > 1 imply in an aggregated pattern, and *I_a_* < 1 in a regular pattern. A two-sided hypothesis test was used to evaluate deviation of *I_a_* = 1, from the null hypothesis of randomness (Perry 1998). The analysis was performed using a local clustering correction (Li et al. 2012) with 2,000 randomizations for each dataset by the SADIE function of the EPIPHY package.

#### Spatiotemporal analyses

Two additional approaches (binary power law and spatial association) were used to quantify the spatiotemporal dynamics of SLB incidence. The binary power law is the relationship between the observed variance of diseased leaves and the theoretical variance of a random distribution (Hughes and Madden 1992). A logarithmic transformation can be used to depict the relationship between datasets by 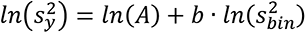, in which 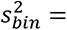 *y*(1 – *y*)/*n* and *A* = *a*·*n^b^*, where *y* is used to estimate the binomial variance, *n* the total number of plants in sampling unit and *a* and *b* are the parameters to be estimated from the data (Madden et al. 2007). When *A* = *b* = 1, the parameters support the binomial distribution on the assumption of randomness. If *A* > *b* = 1, the observed variance is higher than the expected variance and the degree of aggregation is constant. If *A* and *b* > 1, the degree of aggregation is not constant and varies with incidence in a systematic manner. The binary power law was applied for each dataset considering the repeated assessments. The modified Welch’s *t*-test was performed against the null hypothesis of randomness (Welch 1947). The formulas for binary power law analysis were based on (Hughes and Madden 1992; Madden et al. 2018) and performed in R.

In addition to the *I_a_*, *SADIE* measures the contribution of diseased individuals in each sampling unit to a patch (*v_j_*) or a gap (*v_i_*) in the spatial arrangement of SLB incidence. The clustering indices were used to obtain a local spatial association (χ*_k_*) between patches and gaps for each sampling unit (Perry and Dixon 2002). The average of χ*_k_* is the overall spatial association (*Χ*), that is a simple correlation coefficient between the clustering indices of each dataset (Perry and Dixon 2002). A modified *t*-test for spatial association was used to test the significance of *Χ* (Clifford et al. 1989; Dutilleul et al. 1993). Analyses were performed using SADIE and MODIFIED.TTEST functions from the EPIPHY and SPATIALPACK (Osorio et al. 2016) packages, respectively.

### Seed testing

The presence of *S. vesicarium* in commercial onion seedlots not treated with fungicide obtained from 2016 (n = 13) and 2017 (n = 15) cropping seasons was assessed using plate tests. Organic seedlots were chosen to maximize potential for detecting seed transmission of *S. vesicarium*, as seed utilized for broad-acre crops is derived from seed crops which receive fungicides, and seed is often treated with fungicide which may inhibit detection. Onion seedlots tested in 2016 were: cv. Ailsa Craig (pelleted), and non-pelleted seed of cv’s. Bridger, Blush, Cabernet, Cortland, Expression F1, Great Western, New York Early, Patterson, Pontiac, Red Hawk, Red Wing, and Zoey (Johnny’s Selected Seeds, Waterville, ME). Non-pelleted seed of cv’s. Bridger F1, Cabernet F1, Candy F1, Cortland F1, Expression F1, Great Western F1, New York Early, Patterson F1, Pontiac F1, Red Hawk, Red Wing, Scout F1, Sierra Blanca F1, Walla Walla Sweet, Zoey F1 were tested in 2017 (Johnny’s Selected Seeds). Two hundred seeds per seedlot were placed (without surface-sterilization) on 2% water agar (20 g/liter agar; Hardy Diagnostics, Santa Maria, CA) + 0.2 g/liter ampicillin; Sigma-Aldrich, St. Louis, MO) in 9 cm diam. Petri dishes. Ten seeds were evenly spaced on each petri dish and incubated at 20°C with a 10 h photoperiod under fluorescent lights. The entire experiment was repeated in each year and data combined. Each seed was examined for fungal growth and sporulation for identification by morphological characteristics at ×60 magnification after 12 days.

### Effect of infested onion residue on SLB epidemics

One field trial was conducted during each of the 2017 and 2018 cropping seasons at the Research North facility of Cornell AgriTech, Geneva, NY. This farm was chosen due to the absence of likely inoculum sources of *S. vesicarium*, due to geographical isolation from commercial onion crops and a lack of *Allium* spp. production in the last 10 years. Both trials were conducted by establishing plots in fall of the previous season. Each plot received 1.5 kg of infested onion leaf residue (12% moisture content) collected from a commercial onion field following harvest in Elba, NY. Treatments consisted of: (i) onion residue spread across the soil surface; (ii) onion residue spread across the surface followed by burial to a depth of 3 to 5 cm; and (iii) a non-inoculated control (no onion residue amendments). Onion residue was applied to plots on 2 November 2016 and 27 October 2017.

Treatments were arranged in a randomized complete block design with six and four replications in 2017 and 2018, respectively. Individual plots were 4.6 m wide × 7.6 m long in both years. The area between plots (10 m) in both directions was sown with oats on 4 May 2017 and 14 May 2018 for each trial respectively, to form a barrier to reduce the probability of *S. vesicarium* transfer between plots. A 1.2 m wide bed was cultivated with a rotovator through the center of plots and black embossed plastic mulch (Empire Drip Supply, Williamson, NY) laid with subsurface drip tape (Empire Drip Supply) on 17 May 2017 and 24 May 2018 for each trial respectively, in preparation for onion planting.

Onions were planted on 5 May 2017 and 23 April 2018 into 50 cell (80 cm^3^) propagation trays at a depth of 1 cm in potting mix (SunshineMix #8 Fafard-2 (Sungro Horticulture, Agawam, MA), and grown in a glasshouse at temperatures of 21/15°C (light/dark) with a 14 h photoperiod until transplanting by hand on 7 June 2017 and 6 June 2018 at the 3 to 4 leaf stage. Plants were fertilized with Peters Excel (ICL Fertilizers, Dublin, OH) at 130 ppm of nitrogen using a Hozon siphon mixer (Phytotronics, Earth City, MO) twice a week while in the glasshouse. Two rows of onions (90 transplants per plot) were transplanted through the plastic mulch spaced at 10 cm within and 30.5 cm between rows. Onions were irrigated via subsurface trickle irrigation as required to optimize plant health. Nutrients were applied through water-soluble nutrients in the irrigation (Jack’s All Purpose 20:20:20 JR Peters Inc., Allentown, PA) at 21-day intervals. Foliar insecticides were applied in each season for onion thrip control. In 2017, five applications were made consisting of: spirotetramat (Movento; 365.4 ml/ha; Bayer CropScience) on 6 and 18 July; cyantraniliprole + abamectin (Minecto Pro; 730.8 ml/ha; Syngenta) on 1 August; and spinetoram (Radiant; 584.6 ml/ha; Dow AgroSciences) on 14 and 16 August. In 2018, spirotetramat was applied on 9 and 20 July.

SLB intensity was quantified at regular intervals throughout the cropping seasons in each year. On each occasion, one fully expanded onion leaf was detached from each of ten randomly selected plants within each plot. Leaves were placed in individual plastic bags with moist paper towel and incubated at 7 to 10 days at 20°C. Each leaf was examined at ×60 magnification using a stereo microscope to confirm the presence of *S. vesicarium* conidia. Disease incidence (in percentage) was calculated by *y* = (*x*/*N*)×100, where *x* is the number of leaves with SLB symptoms, and *N* total number of leaves. SLB incidence was evaluated at 59, 73, 87, 101, and 115 DAP in 2017. In 2018, SLB incidence was evaluated at 34, 48 and 61 DAP. AUDPS (Simko and Piepho 2012) was calculated in each plot based on SLB incidence in 2017. An insufficient number of assessments (<5) prevented the calculation of AUDPS in the 2018 trial.

#### Data analysis

The effect of residue treatment on SLB incidence (in 2017 and 2018) and AUDPS (in 2017) was analyzed using generalized linear modelling. Data from each trial were analyzed separately. Residue placement (treatment) and replication were evaluated as fixed and random effects, respectively. Means were separated by a Fisher’s protected least significant difference test at *P* = 0.05 (Genstat Version 18; Hemel Hempstead, UK).

## Results

### Spatial and temporal analyses of SLB epidemics

#### Disease intensity

The SLB incidence for all datasets ranged from 0 to 0.99, observed at the initial and last assessments, respectively (Fig. 1A). In 2017 and 2018, the median incidence in the datasets was 0.17 and 0.35, respectively, and significant differences between years were not observed (*P* = 0.629; *data not shown*). The size of lesions ranged from 0.1 to 34.9 cm, from the first to last assessment (Fig. 1B). The average and median values of lesion size in the last assessment was 28.3 and 30.4 cm, respectively (*data not shown*). No significant differences in lesion size were observed between years (*P* = 0.457; *data not shown*).

**Fig. 1.**
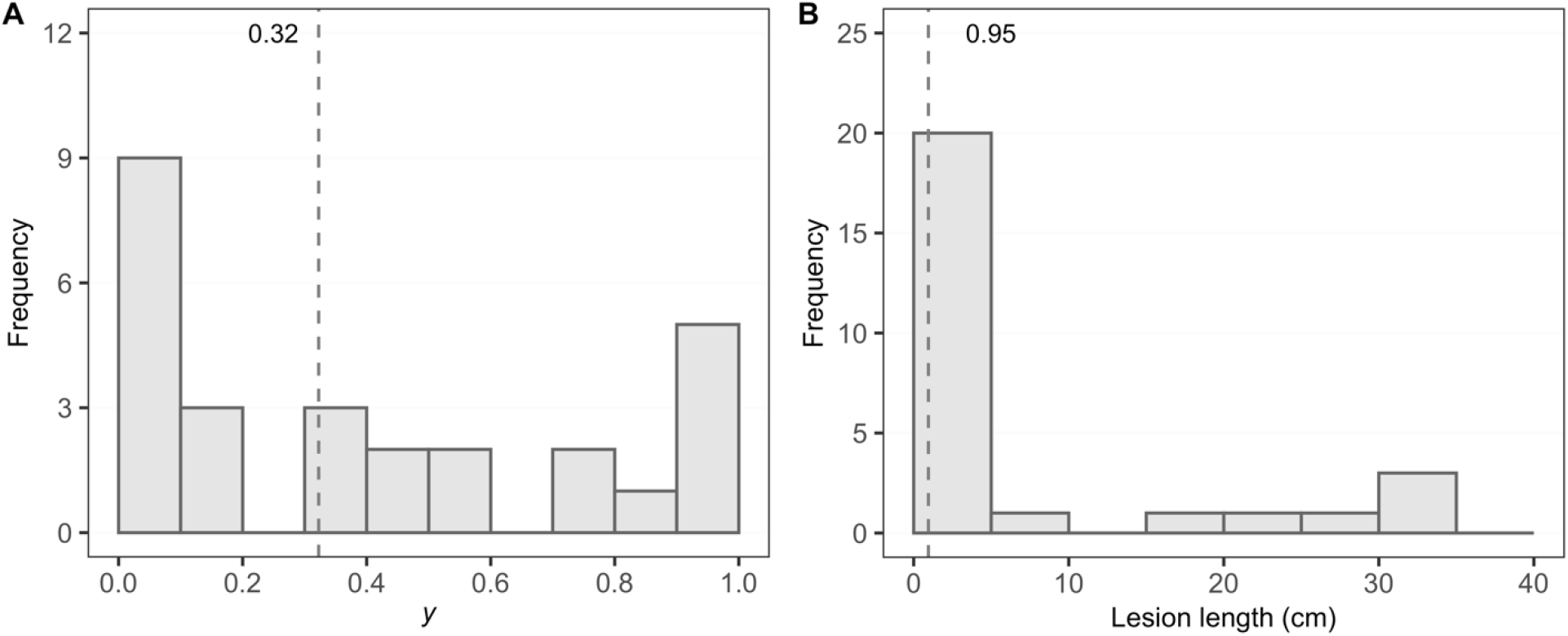
Frequency of datasets containing the **A,** incidence and **B,** size of lesions caused by Stemphylium leaf blight caused by *Stemphylium vesicarium* on onion leaves. Data was collected along three transects per field on each of five assessments in 2017 and four assessments in 2018 from six onions fields in Elba, New York.

#### Disease progress

The temporal change in SLB incidence was a typical sigmoid-type curve in both cropping seasons (Fig. 2). The logistic model provided the best fit to the observed data for 83.3% of the datasets, and resulted in a lower RMSE, larger independence and homogeneity of variances, and *R^2^* compared with the monomolecular and Gompertz models. Based on these parameters, the logistic model was selected for comparisons between the cropping seasons (Fig. 2).

**Fig. 2.**
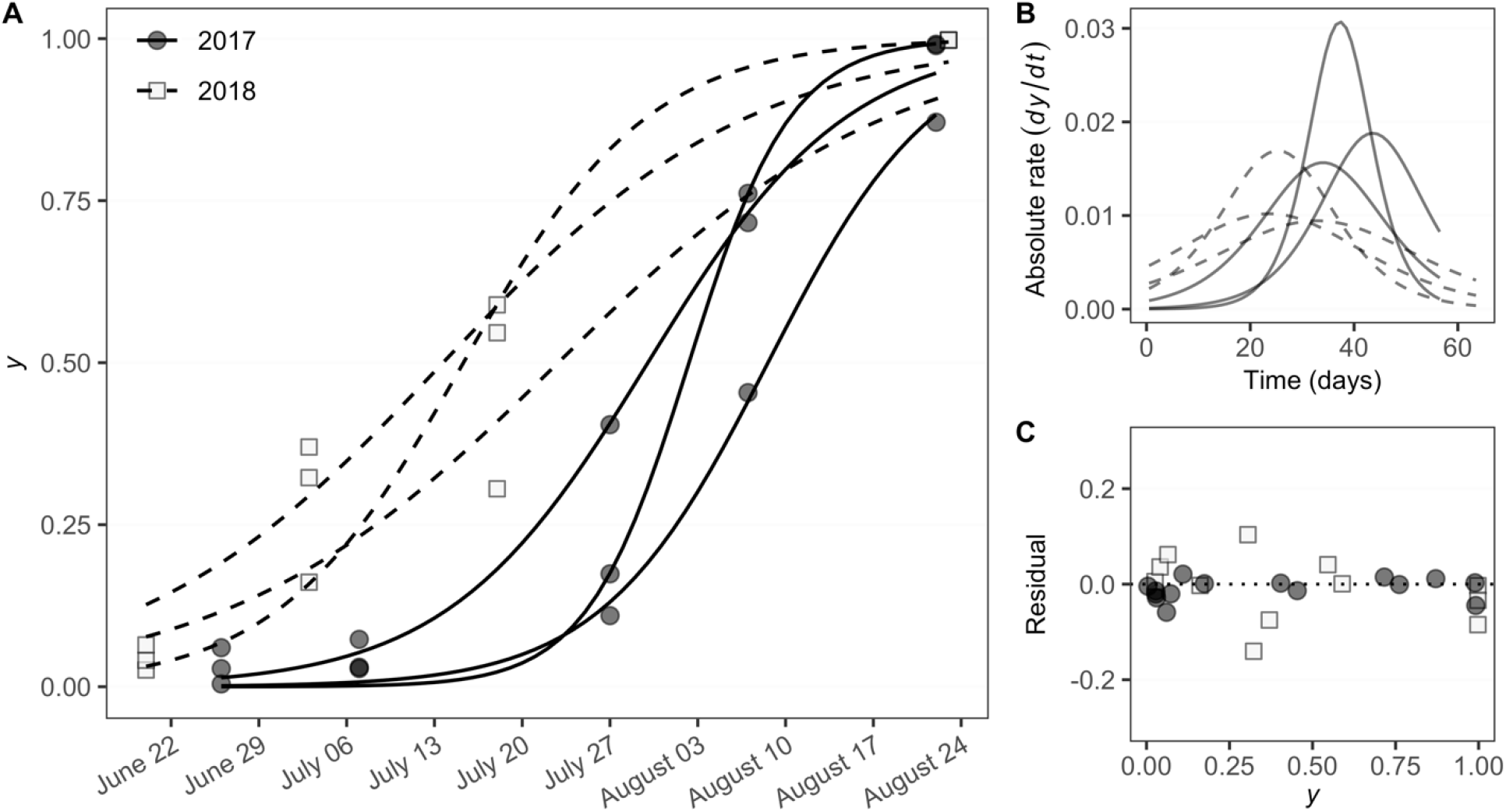
**A,** Stemphylium leaf blight disease progress curves in onion fields in Elba, New York fitted to the logistic model. Three fields were assessed in 2017 (closed points and solid lines) and three fields assessed in 2018 (open squares and dashed lines). **B,** Absolute rate (*dy*/*dt*) based on the logistic model since the first assessment, and **C**, Residual (predicted - observed) *versus* observed incidence using the logistic model.

No difference was observed in initial incidence (*y_0_*; *P* = 0.113), disease progress rate (*r*; *P* = 0.162), time for the epidemic to reach 50% disease incidence (*y_50_*; *P* = 0.063), and final incidence (*y_max_*; *P* = 0.357) between the two cropping seasons (Table 1). The rAUDPS of disease incidence and lesion length were significantly lower (*P* < 0.009) in 2017 than 2018 (Table 1).

**Table 1.** Temporal analyses parameters of logistic model fitted to the disease progress curves of Stemphylium leaf blight epidemics on onion in 2017 and 2018 in Elba, New York

| Season | $y_0^a$ | $r^b$ | $y_{50}$ (days)<sup>c</sup> | $y_{max}^d$ | rAUDPS<sup>e</sup> | |
| --- | --- | --- | --- | --- | --- | --- |
|  |  |  |  |  | Incidence | Lesion length |
| 2017 | 0.005 | 0.174 | 38.4 | 0.95 | 0.004 | 0.059 |
| 2018 | 0.078 | 0.098 | 28.3 | 0.99 | 0.007 | 0.150 |
| *P-value* | 0.113 | 0.162 | 0.063 | 0.357 | 0.008 | 0.009 |

<sup>a</sup> Disease incidence at epidemic initiation.
<sup>b</sup> Epidemic progress rate.
<sup>c</sup> Time for disease incidence to reach 50%.
<sup>d</sup> Disease incidence at final assessment.
<sup>e</sup> Relative Area Under Disease Progress Stairs.

#### Weather data

Significant differences were observed for minimum, maximum and average temperature between the cropping seasons (*P* < 0.002). The temperatures during the 2017 season were, on average, 2°C lower (20.9°C) than in 2018 (22.8°C; Fig 3). Significant differences were not observed (*P* > 0.05) for dewpoint, precipitation, and relative humidity (Fig. 3).

**Fig. 3.**
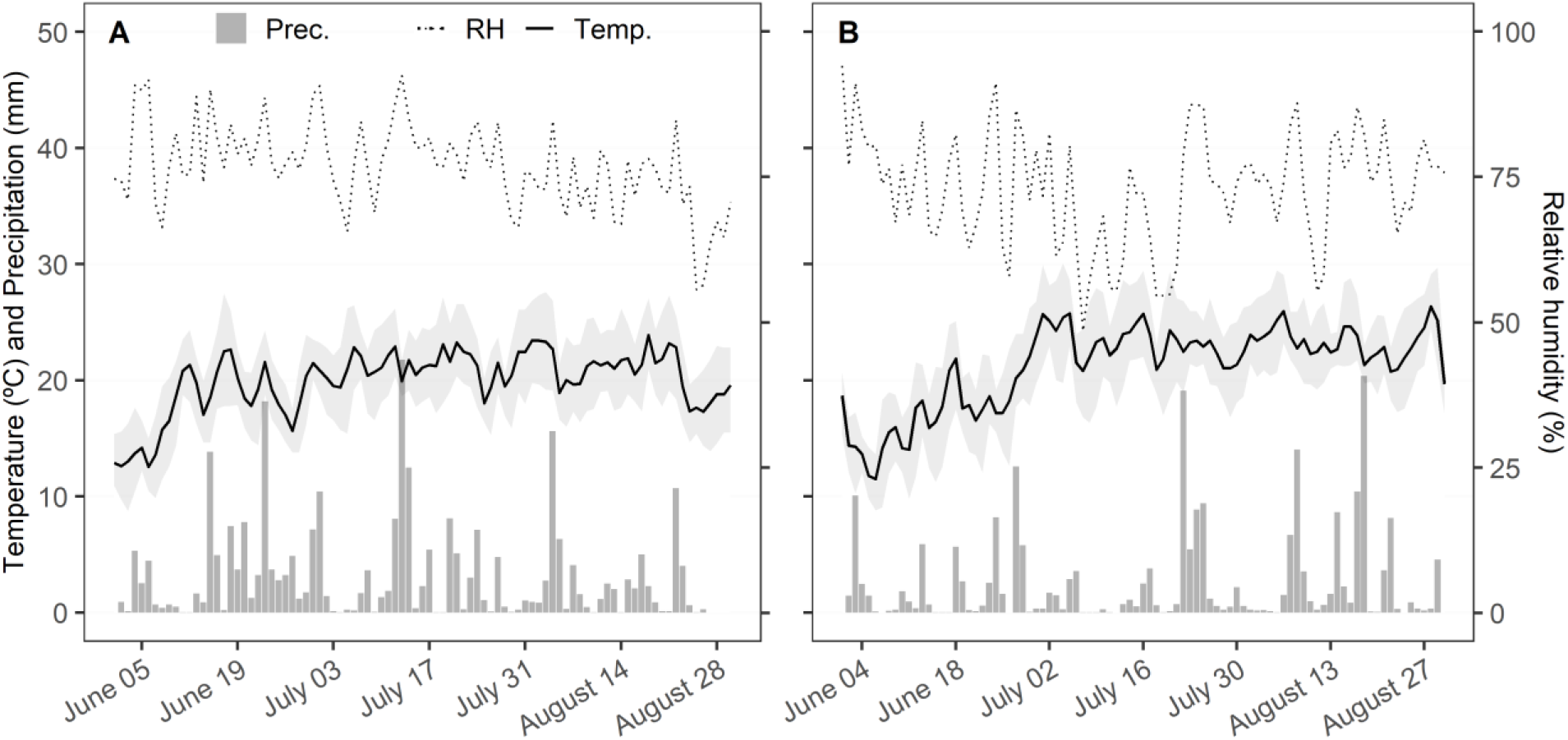
Daily recordings of minimum, maximum (shaded area), and average temperature (Temp.), relative humidity (RH), and precipitation (Prec.) predicted for the centroid of three onion fields in Elba, NY, in **A**, 2017, and **B**, 2018.

#### Spatial analyses

Spatial pattern analyses were performed using all datasets. The binomial distribution was adequately calculated for all datasets and fit (*P* < 0.05) was observed for 63.9%. The beta-binomial distribution was adequately calculated for 67.7% of the datasets by maximum likelihood, from which 94.4% were consistent by the χ^2^ goodness-of-fit test. Values of the beta-binomial aggregation parameter, *θ*, ranged from 0.01 to 0.47 with a median of 0.08 (Fig. 4A) and no significant (*P* = 0.143) difference was observed between seasons (Table 2). However, the LRS test indicated that disease incidence was best described by the beta-binomial than binomial distribution in only half (50%) of datasets (Fig. 4A). Significant differences in the proportion of the fields described by beta- or binomial distributions were not observed between years (Table 2).

**Fig. 4.**
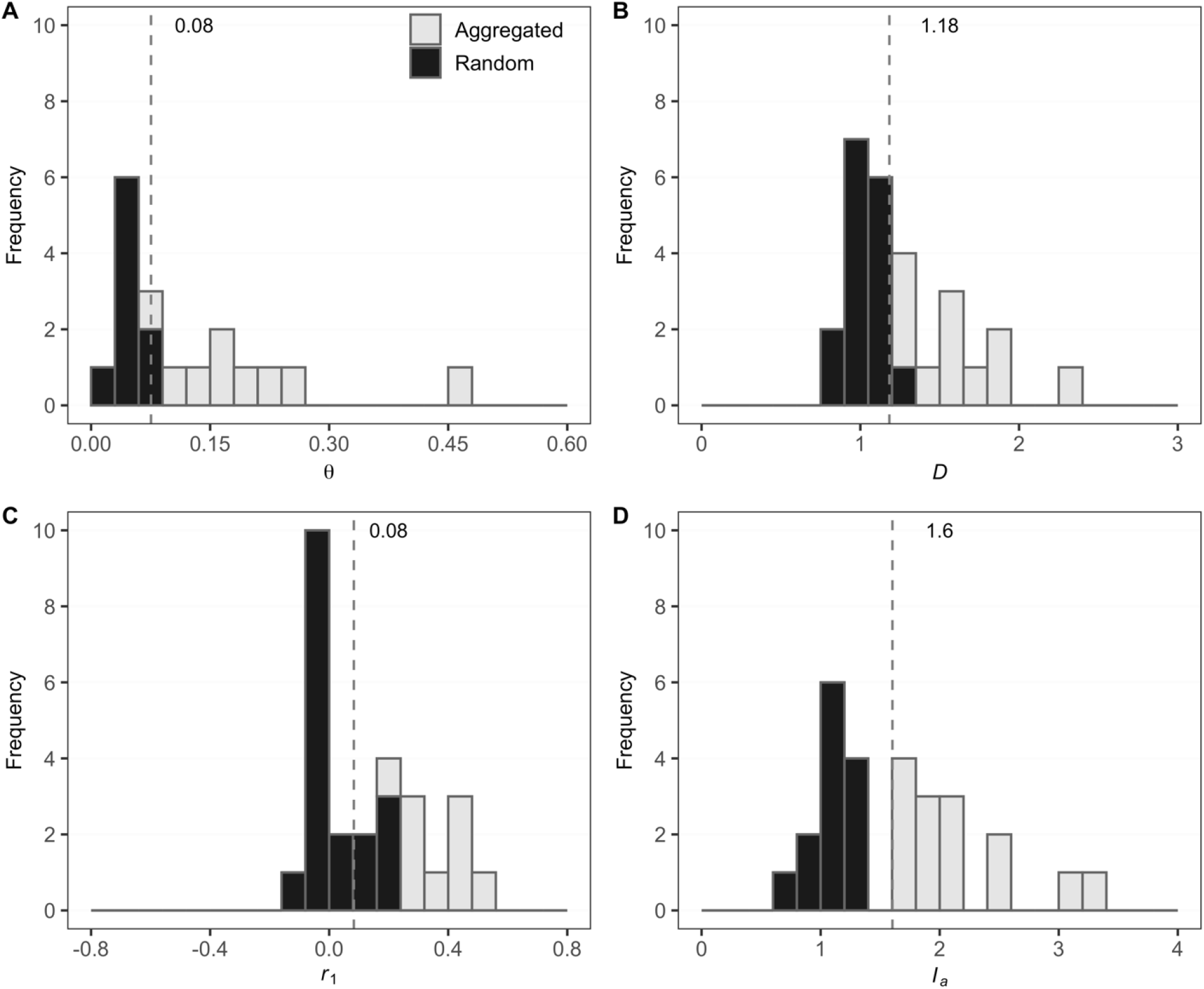
Statistics describing the spatial attributes of Stemphylium leaf blight (SLB) epidemics caused by *Stemphylium vesicarium* in onion fields in Elba, New York, in 2017 and 2018. **A,** Frequency of the aggregation parameter (*θ*) of the beta-binomial distribution, **B**, index of dispersion (*D*), **C**, autocorrelation function on the first spatial lag (*r_1_*), and **D**, index of aggregation (*I_a_*) of Spatial Analyses by Distance IndicEs (*SADIE*). Median values are represented by dashed lines. The classification of datasets as aggregated or random patterns is based on statistical tests (*P* = 0.05).

**Table 2.**
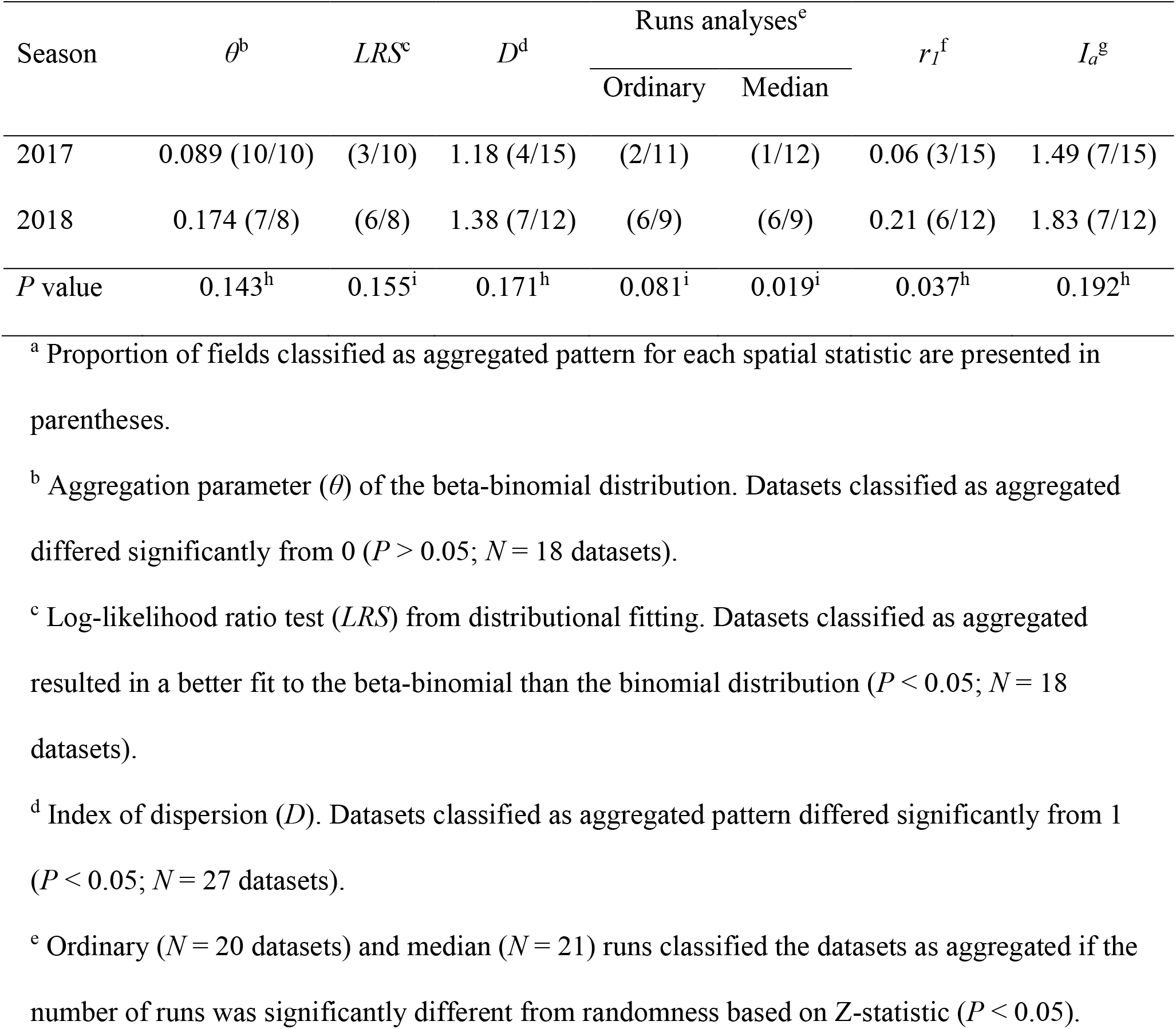
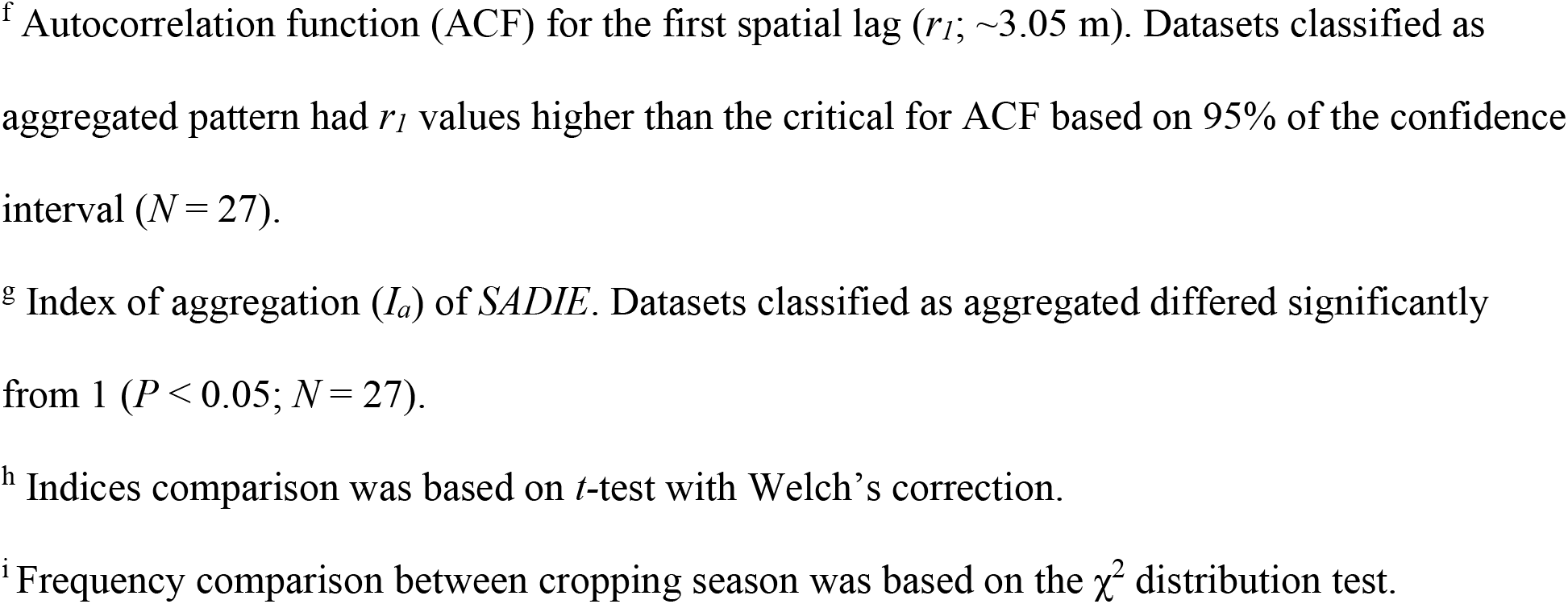
Average values and proportion of datasets classified as having aggregated patterns according to distribution fitting, the log-likelihood ratio test, index of dispersion, ordinary and median runs analyses, spatial autocorrelation, and Spatial Analyses by Distance IndicEs (*SADIE*) according to Stemphylium leaf blight incidence in onion fields in Elba, New York in 2017 and 2018^a^

| Season | $\theta^b$ | *LRS*<sup>c</sup> | *D*<sup>d</sup> | Runs analyses<sup>e</sup> | | $r_I^f$ | $I_a^g$ |
| --- | --- | --- | --- | --- | --- | --- | --- |
|  |  |  |  | Ordinary | Median |  |  |
| 2017 | 0.089 (10/10) | (3/10) | 1.18 (4/15) | (2/11) | (1/12) | 0.06 (3/15) | 1.49 (7/15) |
| 2018 | 0.174 (7/8) | (6/8) | 1.38 (7/12) | (6/9) | (6/9) | 0.21 (6/12) | 1.83 (7/12) |
| *P* value | 0.143<sup>h</sup> | 0.155<sup>i</sup> | 0.171<sup>h</sup> | 0.081<sup>i</sup> | 0.019<sup>i</sup> | 0.037<sup>h</sup> | 0.192<sup>h</sup> |

<sup>a</sup> Proportion of fields classified as aggregated pattern for each spatial statistic are presented in parentheses.
<sup>b</sup> Aggregation parameter ( $\theta$ ) of the beta-binomial distribution. Datasets classified as aggregated differed significantly from 0 ( $P > 0.05$ ; $N = 18$ datasets).
<sup>c</sup> Log-likelihood ratio test (*LRS*) from distributional fitting. Datasets classified as aggregated resulted in a better fit to the beta-binomial than the binomial distribution ( $P < 0.05$ ; $N = 18$ datasets).
<sup>d</sup> Index of dispersion (*D*). Datasets classified as aggregated pattern differed significantly from 1 ( $P < 0.05$ ; $N = 27$ datasets).
<sup>e</sup> Ordinary ( $N = 20$ datasets) and median ( $N = 21$ ) runs classified the datasets as aggregated if the number of runs was significantly different from randomness based on Z-statistic ( $P < 0.05$ ).
<sup>f</sup> Autocorrelation function (ACF) for the first spatial lag ( $r_I$ ; ~3.05 m). Datasets classified as aggregated pattern had $r_I$ values higher than the critical for ACF based on 95% of the confidence interval ( $N = 27$ ).
<sup>g</sup> Index of aggregation ( $I_a$ ) of *SADIE*. Datasets classified as aggregated differed significantly from 1 ( $P < 0.05$ ; $N = 27$ ).
<sup>h</sup> Indices comparison was based on $t$ -test with Welch's correction.
<sup>i</sup> Frequency comparison between cropping season was based on the $\chi^2$ distribution test.

The index of dispersion (*D*) ranged from 0.75 to 2.31 with a median of 1.18. Only 41% of the datasets resulted in aggregation, in which *D* was significantly (*P* < 0.05) higher than 1 (Fig. 4B). In addition, differences were not observed (*P* = 0.171) in *D* values between the cropping seasons (Table 2).

Ordinary runs analysis was adequately computed for 74.1% of the datasets. Aggregation was detected in 40% of the datasets. No significant differences (*P* = 0.081) were observed in the frequencies of the spatial patterns between the seasons by the χ^2^ test. Median runs were adequately calculated for 77.8% of the datasets. Only seven datasets (25.9%) resulted in an aggregated pattern, and from those, six (85.7%), were observed in 2018. Significant differences (*P* = 0.019) were observed in the frequency of aggregated datasets between the seasons (Table 2).

Statistically significant spatial autocorrelation was detected in nine datasets (33.3%) on the first spatial lag (*r_1_*; Fig. 4C) which correspond to 3.05 m. The *r_1_* autocorrelation values ranged from - 0.13 to 0.49 with a median of 0.08 with a weak significant difference (*P* = 0.037) between seasons (Table 2). In the second spatial lag (*r_2_*), autocorrelation values were above CI 95% in nine datasets (*data not shown*). Significant autocorrelation was only detected at the fifth spatial lag (*r_5_*) in four of the datasets. Moreover, in three of these datasets, significant spatial lags at one to four were not detected, suggesting they were randomly above the 95% critical threshold (*data not shown*).

The index of aggregation by *SADIE* (*I_a_*) ranged from 0.744 to 3.34 with a median value of 1.6 (Fig. 4D). The aggregated pattern was significantly (*P* < 0.05) detected in 51.8% of the datasets. No significant differences were observed in *I_a_* between cropping seasons (*P* = 0.192).

#### Spatiotemporal analyses

The incidence data of sequential assessments at the field level were used for spatiotemporal analysis. The relationship between the logarithm of observed and expected variances of the datasets over time was well described by the binary power law (*R*^2^ = 0.987; Fig. 5). The binary power law parameters, ln(*A*) (0.675 ± 0.125) and *b* (1.102 ± 0.026), differed significantly (*P* < 0.001) from 0 and 1, respectively. Larger parameters are suggestive that SLB epidemics exhibited an aggregated pattern, and the level of aggregation varied with disease incidence. Significant differences between the cropping seasons were not detected (*P* ≥ 0.182) for both binary power law parameters (Fig. 5). Individual analyses at field level allowed us to quantify the changes in the aggregation level over time. Coefficients of determination were larger than 0.954, and ln(*A*) ranged from 0.367 to 1.128 with a median of 0.743, and *b* from 1.054 to 1.204 with a median of 1.11. For these set of data, 83.3% had ln(*A*) > 0, suggestive of aggregation, and 66.7% had *b* > 1, meaning the level of aggregation varied with disease incidence (*data not shown*). For spatial association, *SADIE*’s clustering indices were separated into gaps (*v_i_*) and patches (*v_j_*), and ranged from -3.75 to -0.62, and from 0.81 to 3.63, with median values of -1.24 and 1.59, respectively. Differences between the years were not detected (*P* = 0.191). Spatial associations were detected in only one pair of successive assessments (7 July and 27 July) for one dataset (*P* = 0.034) collected in 2017. In the other datasets and pairs of assessments, the *X* values ranged from -0.31 to 0.57 with a median of 0.11 (Table 3). No significant difference (*P* = 0.112) was observed between the cropping seasons.

**Fig. 5.**
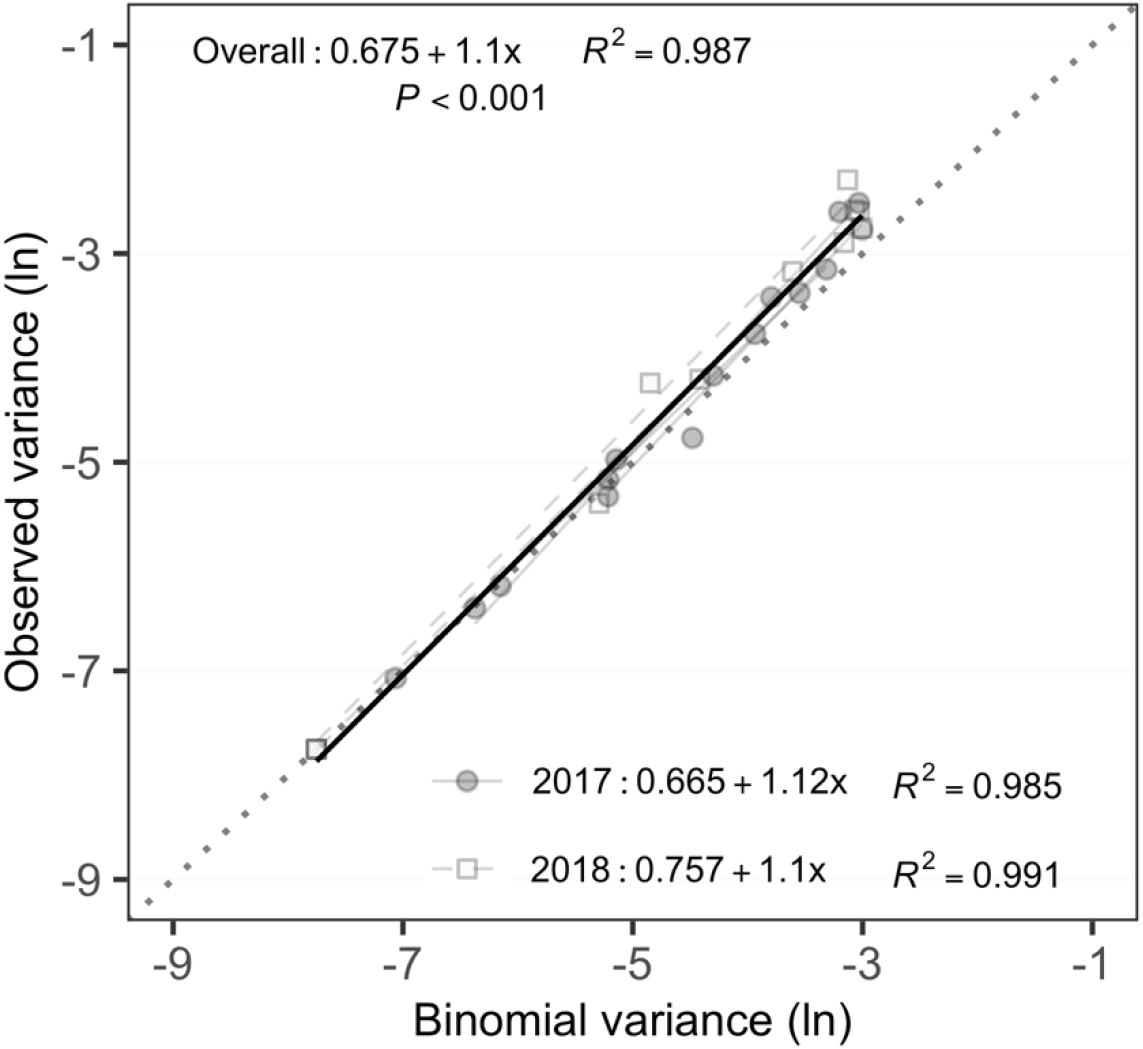
Relationship between logarithms of the observed and binomial variance of the incidence of onions leaves with Stemphylium leaf blight caused by *Stemphylium vesicarium* in Elba, New York in 2017 and 2018. Overall linear regression for six fields (black solid line), and for individual fields (gray lines) assessed in 2017 (closed circles) and 2018 (open squares). The binomial (1:1) line is represented by the dotted line. The linear regression for binary power law parameters is presented. Significant differences were not observed for binary power law parameters, ln(*A*) (*P* = 0.182) and *b* (*P* = 0.243), between cropping seasons.

**Table 3.** Local clustering indices obtained by SADIE and overall spatial association (*X*) for two successive assessments (second and third) of Stemphylium leaf blight incidence in onion fields in Elba, New York, in 2017 and 2018

| Season | Assessment | Clustering indices<sup>a</sup> |  | Overall association, |
| --- | --- | --- | --- | --- |
| | | $v_i$ | $v_j$ | $X^b$ |
| 2017 | 7 Jul | $-0.95 \pm 0.26$ | $1.05 \pm 0.25$ | $-0.142 \pm 0.14$ |
| | 28 Jul | $-2.81 \pm 0.55$ | $2.81 \pm 0.84$ | |
| 2018 | 3 Jul | $-1.28 \pm 0.26$ | $1.59 \pm 0.44$ | $0.284 \pm 0.26$ |
| | 18 Jul | $-1.80 \pm 0.88$ | $1.85 \pm 0.78$ | |

<sup>a</sup> Patch ( $v_j$ ) and gap ( $v_i$ ) clustering indices.
<sup>b</sup> Overall association ( $X$ ) and standard deviation.

### Seed testing

*Stemphylium vesicarium* was not observed on seed of any of the 13 or 15 onion cultivars from the selected commercial onion seedlots in 2016 and 2017, respectively. Other fungi observed growing from seed at low incidences (< 5%) included *Cladosporium* spp., *Fusarium* spp., *Penicillium* spp., *Ulocladium atrum*, and an unidentified Zygomycete.

### Effect of infested onion residue on SLB epidemics

In 2017, residue treatment had no significant effect on SLB incidence at each of the individual assessments (*P* ≥ 0.086) except for 87 DAP (*P* = 0.015). *S. vesicarium* was first detected in plots of surface and buried treatments at 59 DAP but was not detected in plots with no residue until 87 DAP. Significant treatment effects were observed on SLB incidence and epidemic progression as depicted by AUDPS (*P* = 0.037; Table 4). At 87 DAP, the incidence of SLB was not significantly different in plots receiving residue placed on the surface or buried, but significantly lower in plots without residue compared to those with buried residue. AUDPS was 33.7 to 41.6% lower in plots receiving onion residue compared to those without residue and was not significantly different between plots in which residue was buried and those not receiving residue (Table 4). Similar trends were also observed in the trial conducted in 2018 (Table 5). For example, SLB incidence was significantly lower in plots not receiving residue compared to those with residue regardless of placement on the surface or burial, which in turn were not significantly different from each other at 34 and 48 DAP. However, by the final assessment at 61 DAP, SLB incidence was not significantly different between treatments (Table 5).

**Table 4.** Effect of infested onion residue on the incidence of Stemphylium leaf blight caused by *Stemphylium vesicarium* in onion during subsequent season in small-plot, replicated trials conducted at Geneva, New York in 2017

| Treatment<sup>a</sup> | SLB incidence (%) |  |  |  |  |  |
| --- | --- | --- | --- | --- | --- | --- |
|  | 59 DAP<sup>b</sup> | 73 DAP | 87 DAP | 101 DAP | 115 DAP | AUDPS<sup>c</sup> |
| Surface | 8.3 | 5 | 15 ab | 56.7 | 93.3 | 2357 a |
| Buried | 3.3 | 3.3 | 25 a | 33.3 | 86.7 | 2077 ab |
| No residue | 0 | 0 | 1.7 b | 26.7 | 76.7 | 1377 b |
| LSD<sup>d</sup> | - | - | 14.6 | - | - | 735.2 |
| *P-value* | 0.216 | 0.178 | 0.016 | 0.086 | 0.312 | 0.037 |

<sup>a</sup> Means followed by the same letter are not significantly different.
<sup>b</sup> DAP = Days After Planting.
<sup>c</sup> AUDPS = Area Under the Disease Progress Stairs (measure of epidemic progress). Larger values indicative of higher disease severity and faster spread.
<sup>d</sup> LSD = Least Significant Difference ( $\alpha = 0.05$ ); na = not applicable when $P > 0.05$ .

**Table 5.** Effect of infested onion residue on the incidence and severity of Stemphylium leaf blight caused by *Stemphylium vesicarium* in onion during subsequent season in small-plot, replicated trials conducted at Geneva, New York in 2018

| Treatment<sup>a</sup> | Incidence (%) |  |  | Lesion length |  |  | *S. vesicarium* sporulation (%) |  |  |
| --- | --- | --- | --- | --- | --- | --- | --- | --- | --- |
|  | 34 DAP<sup>b</sup> | 48 DAP | 61 DAP | 34 DAP | 48 DAP | 61 DAP | 34 DAP | 48 DAP | 61 DAP |
| Surface | 50 a | 70 a | 45 | 0.8 a | 2.3 a | 1.9 a | 6.4 a | 8.6 | 7.3 a |
| Buried | 37.5 a | 50 a | 40 | 0.5 ab | 1.4 b | 1.7 a | 3.8 ab | 5.9 | 5.6 b |
| No residue | 12.5 b | 17.5 b | 22.5 | 0.3 b | 0.7 b | 1.1 b | 1.9 b | 3.4 | 2.3 c |
| LSD<sup>c</sup> | 16.6 | 25 | - | 0.4 | 0.8 | 0.4 | 3.4 | - | 1.8 |
| *P-value* | 0.004 | 0.006 | 0.131 | 0.029 | 0.011 | 0.013 | 0.049 | 0.058 | 0.002 |

<sup>a</sup> Means followed by the same letter are not significantly different.
<sup>b</sup> DAP = Days After Planting.
<sup>c</sup> LSD = Least Significant Difference ( $\alpha = 0.05$ ); na = not applicable when $P > 0.05$ .

## Discussion

The results from this research have provided new quantitative information surrounding the relative importance of various inoculum sources on SLB epidemics in large-scale muck onion fields in NY. Infested seed was hypothesized as playing a role in the SLB epidemics in onion fields in 1990 (Lorbeer 1993) and *S. vesicarium* can be seed transmitted in onion (Aveling 1993; Aveling et al. 1993). In this study, the lack of detection of *S. vesicarium* in 28 untreated commercial onion seed lots suggests that this path may not currently be a significant means of primary inoculum introduction in this pathosystem. Seed destined for the establishment of conventional onion fields is often grown in climatic regions not conducive to SLB, seed crops are treated with fungicides, and the seed itself is often also treated with fungicides, factors which are likely to reduce the potential for seed transmission. The random spatial distributions of SLB epidemics in most of the datasets collected in this study are therefore unlikely to be due to the introduction of *S. vesicarium* on infested seed. However, further studies inspecting onion seeds used specifically in large-scale NY fields are required to confirm this finding. In this study, the health status of transplants arriving from outside NY for planting was not evaluated. Transplants have been identified as a significant source of IYSV and viruliferous thrips for direct-seeded crops in the Elba muck region (Leach et al. 2018). Transplants may also have the potential to act as an inoculum source for SLB either by introducing inoculum following infections at other locations or providing an early crop for SLB to establish and subsequently provide inoculum for direct seeded crops in the region. In a preliminary study of an onion crop established by transplants in Elba, NY in late May, *S. vesicarium* was present in 90% of leaves with senescent tips at a time when direct seeded crops were establishing (F. S. Hay, *unpublished data*). Similarly, SLB has been detected on senescent leaf tips of volunteer onions early in the spring. Further exploration into the importance of SLB colonization of senescent leaf tips of transplants and volunteer overwintering onions in the SLB epidemic is warranted.

In 2015, *S. vesicarium* was the most frequent (> 85%) pathogen associated with diseased leaves of onions in organic and conventional farms in NY (Pethybridge et al. 2016). Similar trends were observed in both years of this study, 2017 and 2018, with a high incidence of SLB and rapid disease progress. Hay et al. (2019) attributed the re-emergence of SLB as an important disease of onion in NY at least in part to the high prevalence of QoI resistant *S. vesicarium* isolates. The SLB epidemics began in late June in both, 2017 and 2018, cropping seasons, and reached 100% incidence in the latter part of the growing season (late August) for most (5/6) of the fields monitored. The logistic model provided the best fit to the change in SLB incidence with time for most epidemics (5/6). The logistic model best describes polycyclic diseases, in which there are multiple disease cycles within a growing season (Madden 1980; Nutter 1997). Therefore, based on the rapid increase in SLB incidence intensity between late June and August, this is most likely the critical infection period for onion due by *S. vesicarium,* and hence when management strategies should be targeted. Reducing the rate of disease progress during this time may lessen the impact of SLB on crop development including improved photosynthetic area resulting in higher yields.

Disease management during this period may not only slow the rate of disease development, but also improve plant development to reduce defoliation, facilitate lodging, and reduce the incidence of post-harvest issues including sprouting and bacterial bulb rots. In addition, the rapid increase in SLB incidence in late June coincides with the increase in temperature in both seasons. The 2018 season was hotter and drier in comparison to 2017 (Fig. 3). The higher AUDPS for incidence and lesion length in 2018 may have arisen from a combination of heat stress and high thrips numbers in crops in 2018, which may have predisposed plants to SLB (C. Hoepting *personal communication*). In greenhouse, trials thrips have been shown to vector conidia of *S. vesicarium*, and thrips feeding increased fungal colonization of leaves and leaf dieback (Leach et al. 2019). Further, in 2018 season, fungicide programs may have been less effective due to the development of resistance to FRAC 7 fungicides (Hay et al., 2019). Other potential modes of inoculum spread that could have contributed to the mostly random spatial distributions of SLB include wind-borne ascospores (Simmons 1969; Basallote-Ureba et al. 1999; Gossen et al. 2021; Katoch and Kumar 2017). The contribution of ascospores to SLB epidemics on onion in NY is poorly understood. In other countries, pseudothecia typically form at the end of the growing season on leaves and stalks (Raghavendra Rao and Pavgi 1975; Basallote-Ureba et al. 1999; Gossen et al. 2021). Abundant, immature pseudothecia are commonly observed on onion leaves later in the growing season in NY (Hay et al. 2019), and ascospores have been observed in pseudothecia on leaves kept in plastic bags in cool storage (5°C) for several weeks (F. S. Hay, *unpublished data*). *Pleospora allii* ascospores released from garlic leaf residues coincides with frequent precipitation, temperatures between 10 to 21°C and low vapor pressure deficit (Prados-Ligero et al. 2003). Similarly, pseudothecia on pear residue develop slowly, but only at high relative humidity (>98%) and with optimum temperatures between 10 and 15°C (Llorente and Montesinos 2004). From the initial lesions resulting from ascospores, conidia are produced, dispersed, and can infect new tissues and plants in the same cropping season, giving rise to the secondary cycle (Gossen et al. 2021). However, the aggregation of diseased leaves typically observed with rain-dispersed, short range pathogens were not detected at the spatial scale sampled in this study.

The lack of dominance of aggregated spatial patterns also discounted obvious disease gradients from field edges. However, the lack of significant spatiotemporal associations in SLB intensity also suggest the dominance of an external (outside the scale of sampling) source of inoculum that may be significant for SLB spread. Thrip-mediated conidial transfer identified by Leach et al. (2020), could also explain the dominance of random patterns of SLB epidemics in onion fields. Moreover, thrip-damaged plants which are also likely to be randomly distributed may also be more susceptible to infection by *S. vesicarium* (Leach et al. 2020). Alternative hosts, not appraised yet, including barley, and volunteer onions, could act as a randomly distributed source for inoculum in the fields. Barley is used as a nurse crop in the establishment of direct seeded onion fields for wind protection. Volunteer onions derived from the past cropping season are also frequently observed in direct seeded fields. Both hosts may act as a green bridge for inoculum between seasons.

Crop residue is also a likely source of *S. vesicarium* inoculum between seasons in the form of mycelium or pseudothecia (Basallote-Ureba et al. 1999; Rossi and Pattori 2009). In the absence of crop rotation, residue management to reduce the probability of inoculum carry-over between seasons is critically important. In this study, SLB incidence was significantly higher in the subsequent season if residue was left on the surface in fall compared to plots established with no residue. This finding suggests that removal of residue after harvest in fall may have substantial effects for the SLB epidemic in the next season in fields not exposed to external sources of inoculum. Further research should explore additional means of residue management such as deep burial, or the addition of urea or lime sulfur to enhance microbial activity and breakdown of leaf tissue (Beresford et al. 2000; Mondal and Timmer 2003), or mechanical means to increase the surface area of the residue to promote breakdown and incorporation into the soil directly after harvest. Crop rotation to reduce the carry-over of inoculum between seasons is also strongly encouraged. Residue management and crop rotation may be particularly useful in fields which are geographically isolated from external sources of inoculum. However, the predominately random spatial distribution of *S. vesicarium* in onion in our study highlights that inoculum external to the field are important in muck regions of NY. Future research should investigate whether these sources are from adjacent onion fields, or alternative hosts of *S. vesicarium*, and prioritize quantifying the role of the teleomorph, as this stage of the pathogen lifecycle may have a substantial role in the epidemiology of SLB in NY.

## Acknowledgements

This research was supported by the United States Department of Agriculture National Institute of Food and Agriculture (USDA-NIFA) Hatch project NYG-625445, managed by Cornell AgriTech, Cornell University, Geneva, NY, the Federal Capacity Fund Initiative Project 2016-17-149, and the USDA-NIFA Crop Protection and Pest Management Applied Research and Development Program project number 2016-70006-25838. We thank Carol Bowden, Elizabeth Burbine, Sean Murphy, Nicolette Nault, and David Strickland (listed alphabetically by surname), and the field research unit at Geneva, for excellent technical assistance. We are also grateful to the onion growers involved in this study for field access and allowing collection of diseased samples.

## Notes

### Competing Interest Statement

The authors have declared no competing interest.

